# Biological function and mechanism of *flpL* gene in regulating pathogenicity of *Aeromonas hydrophila*

**DOI:** 10.1101/2025.02.17.638635

**Authors:** Hanyang Jiao, Kemei Liu, Ziyi Liao, Adeeba Naseer, Hua Ye, Hao Xu, Yun Li, Yongyao Yu, HuiQing Mei, Ronghua Wu

## Abstract

*Aeromonas hydrophila* is human-animal commensal bacterium that seriously threatens the development of aquaculture and human health. Type IV pili (T4P) are essential for bacterial physiological function. The relationship between the molecular mechanism of *flpL* gene affiliated to T4P and the pathogenicity is still unclear. In the research, *A. hydrophila* with a genetically stable deletion of *flpL* gene (Δ*flpL*) was constructed. The median lethal dose of Δ*flpL* for Crucian carp was 4.87 times higher compared to wild-type strain, suggesting that Δ*flpL* significantly reduced pathogenicity. The attenuation may be attributed to a reduced capacity for biofilm formation and downregulation of the expression levels of virulence-related genes in Δ*flpL*. Meanwhile, the significant increase in swimming capacity and adhesion can be related to the upregulation of *flpC, tapM, tapB* and *tapP* genes. In conclusion, *flpL* gene plays essential role in the pathogenicity, adhesion, motility, and capacity for biofilm formation in *A. hydrophila*. The research aims to clarify the pathogenesis of *A. hydrophila*, and lay the foundation for developing live attenuated vaccines through the targeted modification *flpL* gene.

## 1. Introduction

*Aeromonas hydrophila* is a gram-negative bacterium frequently present in aquatic habitats (Ji *et al*, 2015). It can infect fish, domesticated animals and mammals, causing a critical threat to the aquaculture industry and public security (Fang *et al*, 2004). *A. hydrophila* is capable of causing significant disease outbreaks in various economically essential fish species, including Crucian carp (*Carassius auratus*) (Feng *et al*, 2022), *Micropterus salmoides* (Yang *et al*, 2023) and *Oreochromis niloticus* (Aly *et al*, 2023). The fish infected by *A. hydrophila* usually appear focal symptoms, including skin ulcers, tissue swelling and surface hyperemia, leading to hemorrhagic septicemia and ulcerative syndrome (Fernandez-Bravo & Figueras, 2020). In recent years, there was rising incidence of aquatic animal deaths due to *A. hydrophila* infections (Suresh & Pillai, 2023; Li *et al*, 2023b). These occurrences have resulted in significant economic losses and have severely hindered the advancement of aquaculture. Recently, the prevention and treatment of *A. hydrophila* infections frequently involved the antibiotics (Khalil *et al*, 2021). However, large and frequent usage of antibiotics in aquaculture, not only caused serious pollution to the aquatic environment, but also raised the problem of drug-resistance and residues in fish (Liang *et al*, 2024). Currently, various types of vaccines have emerged to combat *A. hydrophila* infections, including inactivated vaccines (Zhang *et al*, 2020), DNA vaccines (Liu *et al*, 2022), subunit vaccines (Zhang *et al*, 2023) and live attenuated vaccines (Li *et al*, 2021). Among these, the live attenuated vaccines closely resemble the natural growth state and are characterized by low production costs, minimal toxicity, and robust immune induction. (Mohd-Aris *et al*, 2019). Deleted virulence genes responsible for pathogenicity were used to develop live attenuated vaccines (Yi *et al*, 2024; Chi *et al*, 2023).

Virulence factors could increase the possibility of pathogenicity to the host (Sarkar *et al*, 2021). The pathogenic factors of *A. hydrophila* are multifactorial, including pili, flagella, adhesins, cytotoxins, hemolysins and lipases (Rasmussen-Ivey *et al*, 2016). Among them, pili are surface structures of bacterium that consist of repeated subunits which interact either covalently or non-covalently. Pili are essential virulence factors of proteobacteria that significantly contribute to bacterial physiology, including adhesion, biofilm formation and cell motility (Lukaszczyk *et al*, 2019). The classification of *A. hydrophila* pili is based on their assembly pathways, which include type I pili and type IV pili. The type IV pili can be further categorized into three distinct types: Flp pili, Tap pili, and MSHA pili. The tight adherence macromolecular transport system represents an essential type II secretion mechanism. The tad gene functions to encode the components required for the assembly of adhesive Flp pili (Tomich *et al*, 2007). The Flp pili were first reported in *Actinobacillus pleuropneumoniae* (Li *et al*, 2019), the causative agent of locally invasive periodontitis. These pili are essential for colony morphology and biofilm formation, which are important virulence factors. The *flpL* gene is located in the *flpABCDEFGHIJKL* gene cluster, contributing to product Flp pili pseudopilin (Boyd *et al*, 2008). Study have revealed that *flp* mutant strain of *A. pleuropneumoniae* significantly reduced capacity of biofilm formation, cellular adherence, resistance against pathogenicity, and survival in swine blood compared with the wide strain (Li *et al*, 2019). The function and molecular mechanism of the *flpL* gene, which belongs to the Flp pili, remain unclear in *A. hydrophila*. Therefore, we hypothesized that the *flpL* gene might be crucial for the pathogenic mechanisms of *A. hydrophila*.

In the study, the *flpL* gene deletion strain of *A. hydrophila* (Δ*flpL*) was generated through homologous recombination. We compared various parameters between the wild-type strain (WT) and Δ*flpL*. These findings elucidated the effects and underlying mechanisms of the *flpL* gene on the pathogenicity and biological properties of *A. hydrophila*. The study will provide theoretical evidence supporting the potential development of Δ*flpL* as a live attenuated vaccine against *A. hydrophila* infections.

## 2. Materials and methods

### 2.1 Bacteria, plasmid, and cultivation

Table 1 summarizes bacteria and plasmids utilized in this research. The WT was obtained from our laboratory’s previous collection (Wu *et al*, 2020). In addition, the WT was cultivated in BHI medium at a temperature of 28℃. The host *Escherichia coli* SM10λpir bacterium was cultured in LB medium at a temperature of 37℃. In order to attain positive selected strains, the concentrations of antibiotics are ampicillin chloramphenicol. 15% sucrose was used to screen for gene deletion strains.

**Table 1.** Bacteria and plasmids used in the study.

| Bacteria or plasmids | Source or Reference |
| --- | --- |
| *A. hydrophila* | Laboratory stock (Wu *et al*, 2020) |
| $\Delta flpL$ | This study |
| *E. coli* SM10λpir | Laboratory stock (Xiong *et al*, 2024) |
| pRE112 | Laboratory stock (Chi *et al*, 2023) |
| pRE112-$\Delta flpL$ | This study |

### 2.2 Establishment of *A. hydrophila* mutant strain (Δ*flpL*)

The *flpL* gene deletion strain was established by referencing our previously constructed method (Xiong *et al*, 2024; Lu *et al*, 2024a; Long *et al*, 2024; Lu *et al*, 2024b). Firstly, the primers were designed using the comprehensive genome sequence of *A. hydrophila* (NCBI Reference Sequence: NC_021290.1). Secondly, upstream and downstream homology arms were amplified separately. Thirdly, the recombinant fragment was constructed by joining the fragments obtained above using overlap PCR. The fragment was ligated to the enzymatic site of pRE112 plasmid to acquire the pRE112-Δ*flpL*. Afterwards, the pRE112-Δ*flpL* was transformed into *E. coli* SM10λpir, and confirmed successful recombination. Next, the SM10λpir with pRE112-Δ*flpL* recombinant plasmid was reorganized into the WT for conjugation. Finally, by Amp and Cm sensitivity and sucrose resistance, the Δ*flpL* mutant confirmed through PCR with *flpL*-a/*flpL*-d primers and direct DNA sequencing of the mutation sites, were selected. The successfully screen colony was inoculated into BHI medium at a ratio of 1:10 per hour, and samples were taken 12 hours later and validated using *flpL*-a/*flpL*-d primers. All primers used for establishing Δ*flpL* are detailed in Table 2.

**Table 2.**
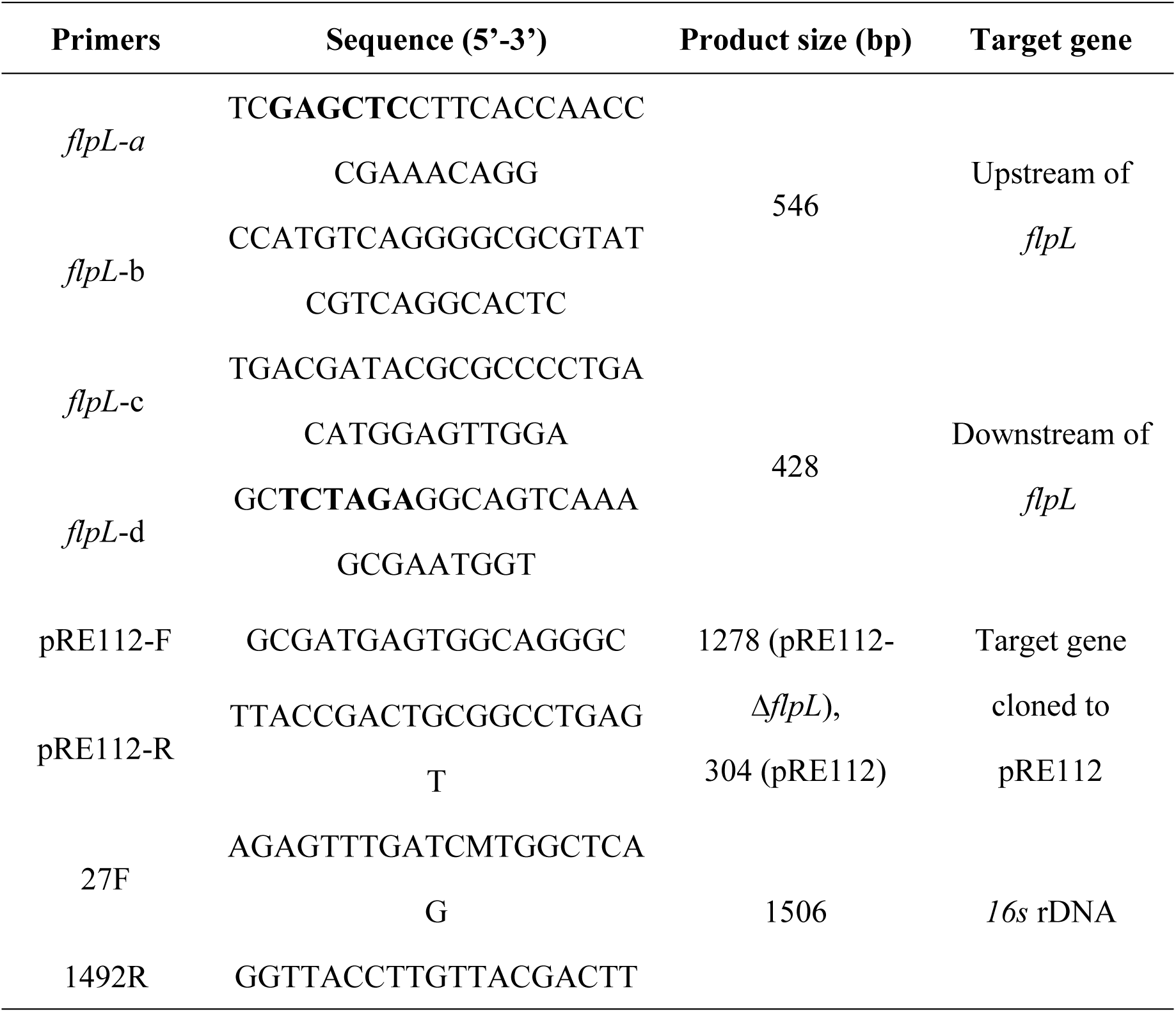
Primers used for establishment of Δ*flpL*.

### 2.3 Median lethal dose (LD_50_)

Healthy Crucian carp, with an average weight of 20 ± 1 g, were sourced from a farm in Chongqing, China. The water temperature was maintained at 26 ± 1℃. During the two-week acclimation period, the fish were fed daily at 1% of their body weight. After confirming the absence of *A. hydrophila* infection through bacteriological testing of eight randomly selected individuals, the experiment proceeded.

To examine the LD_50_ of WT and Δ*flpL*, bacterial suspensions were prepared by diluting the cultures in fresh BHI broth to achieve four different concentrations: 3 × 10^7^ CFU/mL, 3 × 10^6^ CFU/mL, 3 × 10^5^ CFU/mL and 3 × 10^4^ CFU/mL. Crucian carp (n = 12 per group) were randomly assigned to 9 groups. Each group received an intraperitoneal injection of 200 µL from above the four concentration gradients of WT and Δ*flpL*, individually. The control group received an equivalent injection of sterile BHI broth. The fish were monitored for a period of two weeks, during which mortality was recorded following infection until them stabilized. The LD_50_ values of WT and Δ*flpL* for Crucian carp were analyzed according to previous method (Finney, 1985).

### 2.4 Growth curve analyses

To determine the growth capacity, the same concentrations of WT and Δ*flpL* bacterial fluids were respectively inoculated into BHI both. The WT and Δ*flpL* medium were taken every hour, and the optical density (OD) at 600 nm (OD_600_) was monitored until the culture reached a stationary phase. The experiment was replicated at least three times to construct growth curves for both the WT and Δ*flpL* strains.

### 2.5 Hemolytic activity

To measure the haemolytic activity, the same volume of WT and Δ*flpL* was taken during the logarithmic growth phase of bacteria (OD_600_ = 0.5), and then inoculated into Columbia blood agar plates. Afterwards, plates were subsequently cultured at 28℃ for 24 hours to investigate the difference between WT and Δ*flpL*.

### 2.6 Motility assay

The bacterial liquids of WT and Δ*flpL* were respectively aspirated 2 µL to the swimming and swarming nutrient medium (Chi *et al*, 2023), using the same experimental manipulation of hemolytic activity test. This experiment was replicated a minimum of three times to verify swimming and swarming capacity.

### 2.7 Biofilm formation test

The assessment of biofilm formation capacity was based on the methodology that we previously published (Chi *et al*, 2023; Xiong *et al*, 2024). In short, bacterial suspensions of WT and Δ*flpL*, both adjusted to an OD_600_ of 0.5, were inoculated into 96-well plates and incubated for 24 hours. Subsequently, the bacteria in each well were fixed using 99% methanol and stained with 0.1% crystal violet. The dye was then dissolved with 33% glacial acetic acid, and the capacity of biofilm formation between WT and Δ*flpL* was quantified by examining the OD_575_ (Stepanović *et al*, 2000).

### 2.8 Adhesion capacity

Absolute fluorescence quantitative PCR was conducted to determine difference in adherence (Gao *et al*, 2016; Mansour *et al*, 2019). The healthy Crucian carp were randomly allocated into three groups. Afterwards, the fish were subjected to immersion in WT (3 × 10^6^ CFU/mL), Δ*flpL* (3 × 10^6^ CFU/mL) and sterile PBS for a period of 2 hours. Gill tissue samples weighing 30 mg were collected from WT and Δ*flpL* for DNA extraction. Next, the two types of DNA were normalized to the same concentration for qPCR analysis, and Ct values were recorded. The Ct values were brought to the standard curve constructed by our previous study to acquire the *A. hydrophila* of adhered bacteria (Xiong *et al*, 2024). Meanwhile, the gills were uniformly ground and coated on Rimler-shotts medium to observe the difference between WT and Δ*flpL* more clearly.

### 2.9 Expression of virulence-related genes

To reveal the molecular mechanisms with the impact of *flpL* deletion on virulence and biological characteristics, real-time quantitative PCR (RT-qPCR) was conducted to compare the expression levels of virulence-associated genes in WT and Δ*flpL.* Total RNA was extracted from WT and Δ*flpL* strains, followed by the elimination of genomic DNA contamination and synthesis of complementary DNA (cDNA), all performed according to the manufacturer’s protocols using the respective kits. Expression of some essential virulence genes were determined by RT-PCR test, including some type IV pili related genes, *lip*, *aha*, *aerA* and *hly* gene, and analyzed relative expression levels between WT and Δ*flpL*. The dates were analyzed using the 2^-ΔΔCt^ method (Ganger *et al*, 2020). The *16S rRNA* is the internal gene as control. The fluorescence intensities of each relative gene were analyzed in triplicate. Aiming to exclude non-specific amplification problems, and the specificity was validated according to the melting curve of the RT-qPCR products. The experiments were independently performed four times. The primers for RT-qPCR are provided in Table 3.

**Table 3.**
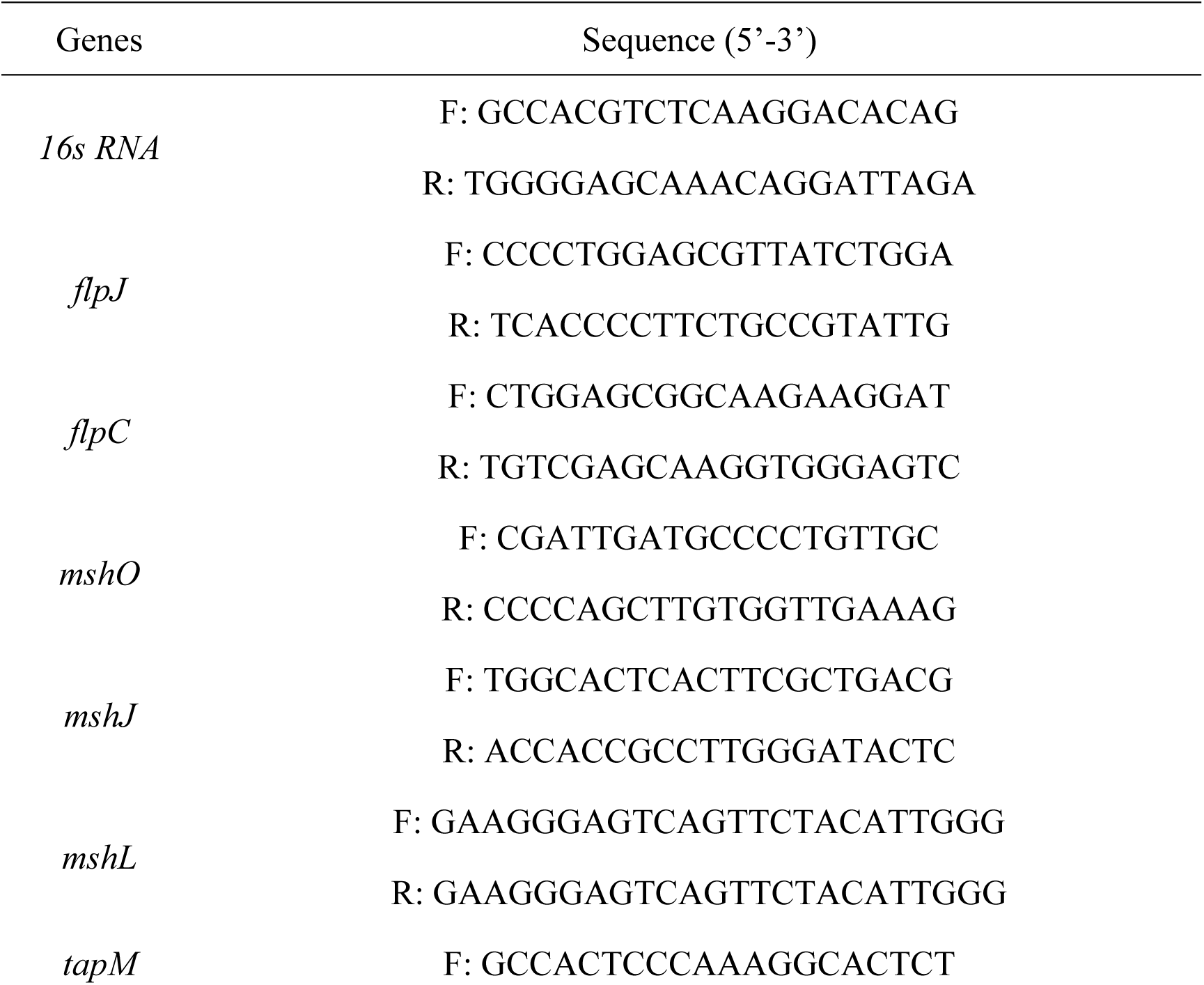
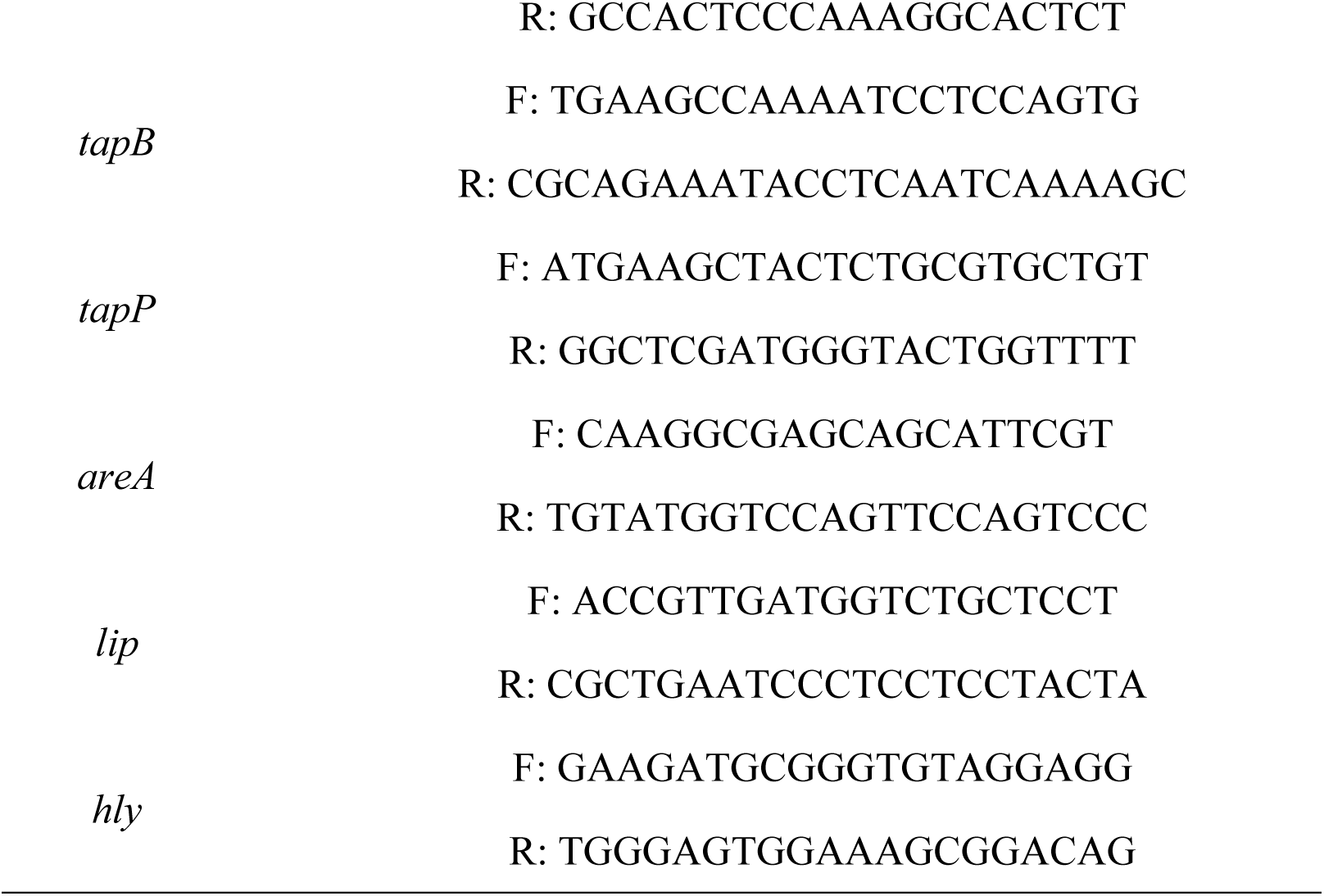
Sequences of the primers for RT-qPCR.

### 2.10 Statistical analysis

Statistical analysis was conducted using SPSS v.26.0 and Origin2021 software. All results are expressed as the mean ± SD (standard deviation), and one-way analysis of variance (ANOVA) was employed for comparisons among groups.

## 3. Results

### 3.1 Establishment of Δ*flpL* strain

The lengths of the homology arms located upstream and downstream of the *flpL* gene were 546 bp and 428 bp, respectively (Fig. 1A). The fusion fragment obtained by overlap PCR, measuring 974 bp (Fig. 1B). The pRE112-Δ*flpL* was successfully transferred into *E. coli* SM10λpir by detecting 1314 bp (Fig. 1C). The first successfully homologous recombination resulted in a double band at 1514 bp and 974 bp (Fig. 1D). The second successfully homologous recombination resulted in a single band at 974 bp (Fig. 1E). Ultimately, the genetic stability assay WT as control was a 1514 bp band and the Δ*flpL* had a 974 bp band. This finding indicated that the Δ*flpL* exhibited stable inheritance and maintain genetic stability (Fig. 1F).

**Fig. 1.**
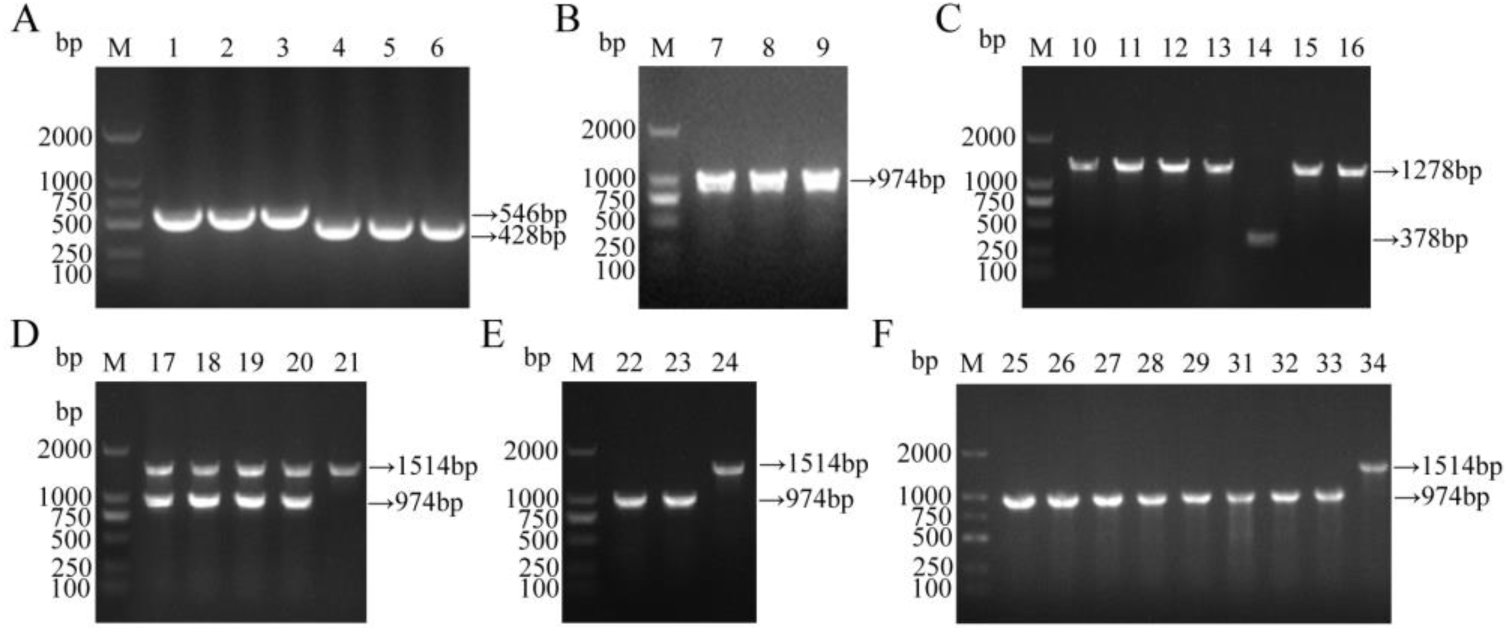
Establishment of Δ*flpL* mutant strain. (A): PCR results for the upstream and downstream homologous arms of *flpL* gene. M is DL2000 DNA Marker, 1-3 are the upstream homologous arms, 4-6 are the downstream homologous arms. (B): Results of upstream and downstream homologous arms overlap PCR product gel recovery. M is DL2000 DNA Marker, 7-9 are the *flpL* gene deletion fragment. (C): Transformation of pRE112-Δ*flpL* recombinant plasmid into SM10λpir verified by colony PCR. M: DL2000 DNA Marker, 10-13 and 15-16 are PCR results of pRE112-Δ*flpL* recombinant plasmid colonies, 14 is SM10λpir harboring the pRE112 plasmid. (D): The results of the first successfully homologous recombinant PCR. M: DL2000 DNA Marker, 17-20 are strains with double bands, 21 is the WT served as the control. (E): The results of the second successfully homologous recombinant PCR. M: DL2000 DNA Marker, 22-23 are strains with targeted bands, 24 is WT served as the control. (F): Results of the genetic stability assessment for the mutant strain. M: DL2000 DNA Marker, 25-33 are randomly sampled Δ*flpL* bacterial samples within 50 generations, 34 is WT served as the control.

### 3.2 LD_50_ values

The LD_50_ values for WT and Δ*flpL* were 5.33 × 10^5^ CFU/tail and 2.60 × 10^6^ CFU/tail, respectively. Following infection with WT, Crucian carp exhibited superficial bleeding and abdominal swelling (Fig. 2A). In contrast, Crucian carp infected with Δ*flpL* showed only superficial hemorrhaging with milder symptoms, with no notable abdominal distension (Fig. 2B). Bacteria obtained from deceased fish were identified by *16S* rDNA universal primers. Subsequent PCR product sequencing confirmed that the isolates were *A. hydrophila*. The results demonstrated that the LD_50_ of Δ*flpL* was 4.87-fold higher than that of WT, suggesting a significant reduction the virulence of *A. hydrophila* (*P* < 0.01).

**Fig. 2.**
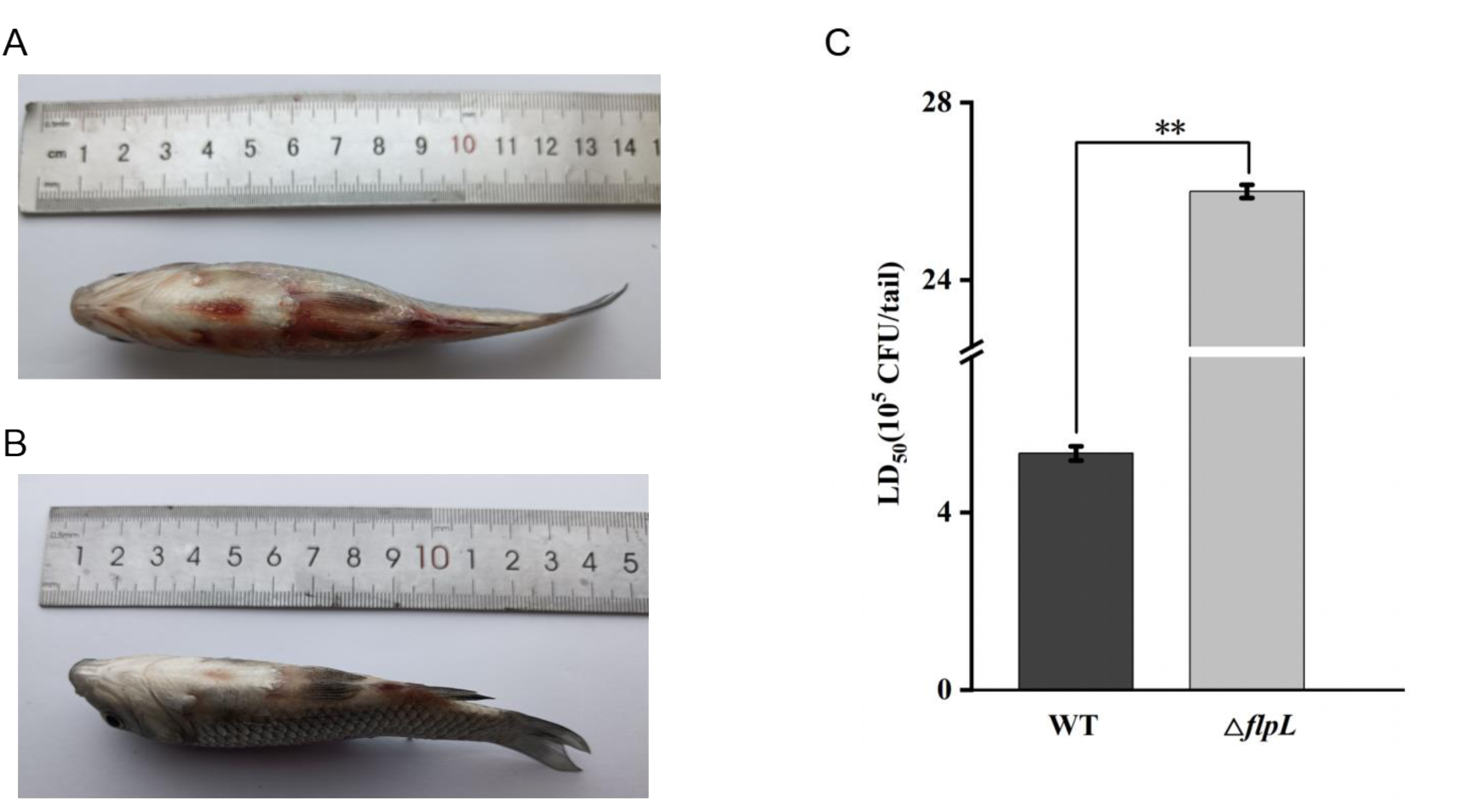
The influence of WT and Δ*flpL* on virulence (**means *P* < 0.01). (A): Symptoms observed in Crucian carp infected with WT. (B): Symptoms obsessed in Crucian carp infected with Δ*flpL*. (C): LD_50_ of WT and Δ*flpL* for Crucian carp.

**Fig. 3.**
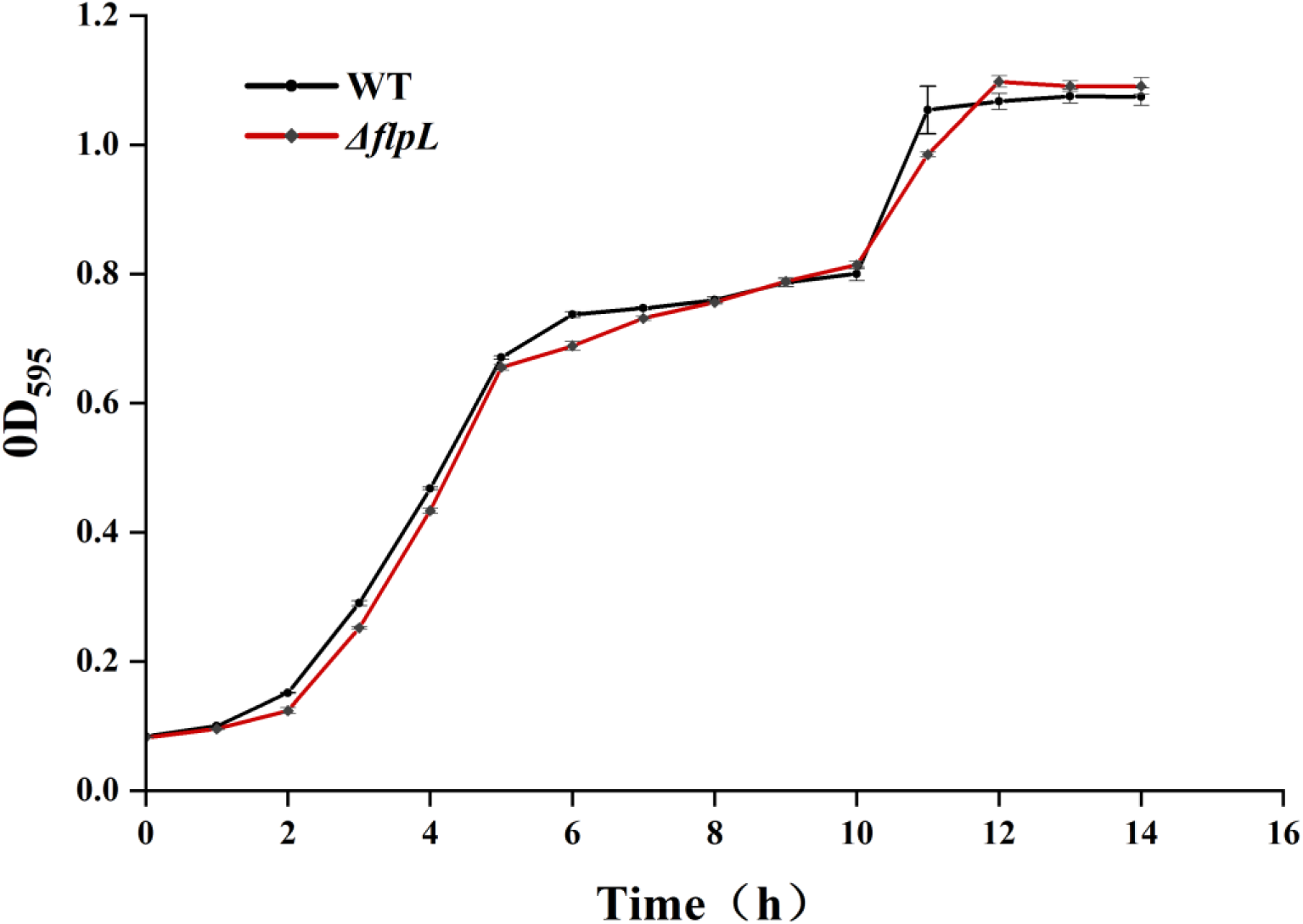
Growth curves of Δ*flpL* and WT.

### 3.3 Influence of the *flpL* gene on the growth capacity

The OD_595_ values for Δ*flpL* and WT did not show significant difference after 15 hours of incubation at 28°C, indicating that deletion of the *flpL* gene did not have a significant influence on the growth capacity.

### 3.4 Effect of *flpL* gene on motility

The diameter of colonies on swimming capacity of Δ*flpL* and WT were 24.59 ± 0.65 mm and 8.98 ± 0.35 mm, respectively (Fig. 4A-B). The results showed that the mutation of the *flpL* gene have a significant enhance of the swimming capacity of *A. hydrophila* (*P* < 0.01). On the swarming plates, the travel distance of Δ*flpL* and WT were 14.09 ± 0.65 mm and 14.91 ± 1.20 mm, respectively. The results demonstrated that there was no significant change on the swarming capacity of Δ*flpL* and WT (*P* > 0.05, Fig. 4C-D).

**Fig. 4.**
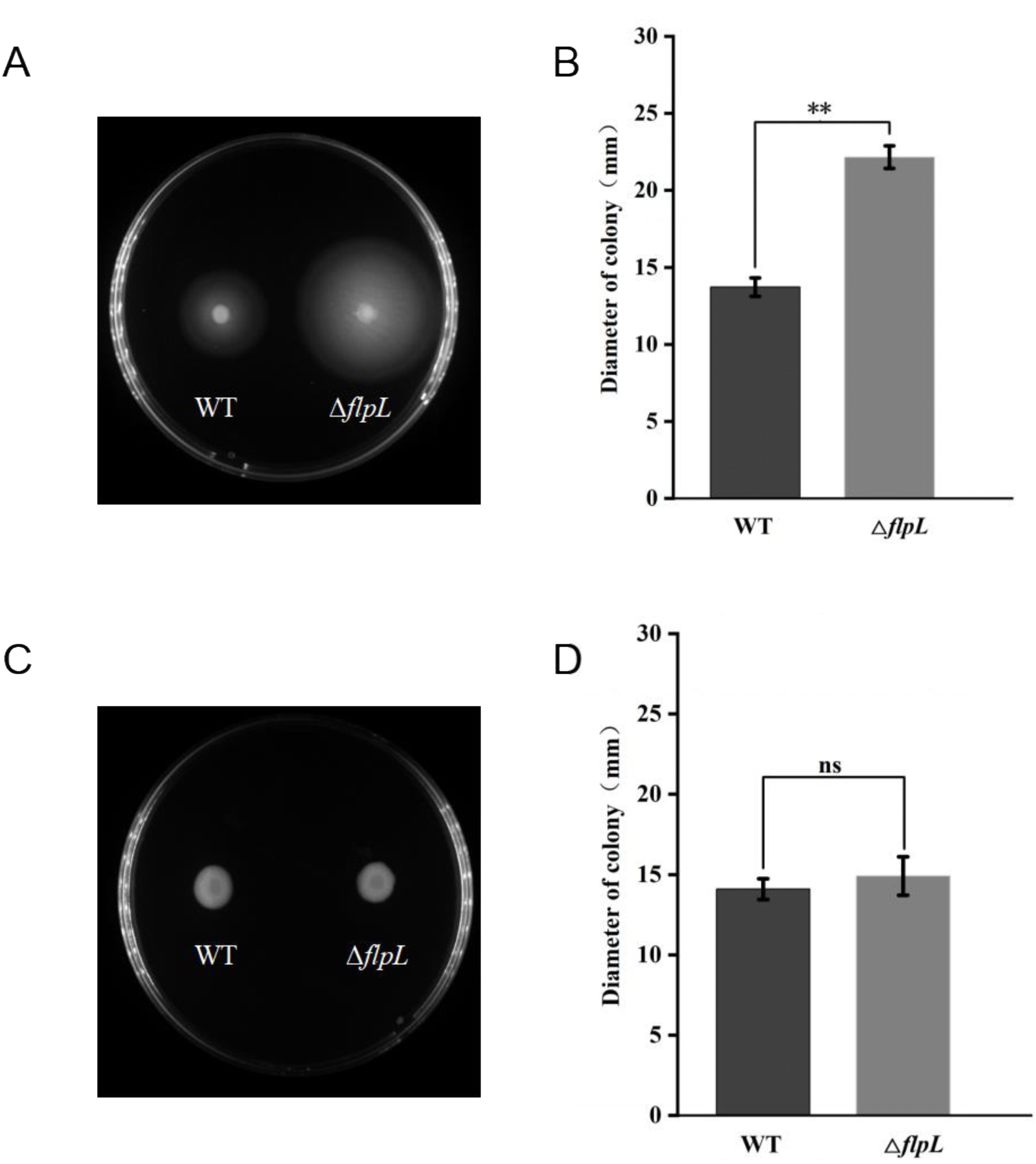
Influence of the *flpL* gene on the motility of *A. hydrophila* (ns represents no significant difference (*P* > 0.05), **means *P* < 0.01). (A, B): Diameter of colonies on swimming capacity of WT and Δ*flpL*. (C, D) Diameter of colonies on swarming capacity of WT and Δ*flpL*.

### 3.5 Hemolytic activity

The result of haemolytic activity demonstrated that both Δ*flpL* and WT exhibited β-hemolysis (Fig. 5A). No significant change in hemolytic activity was observed between Δ*flpL* and WT (*P* > 0.05). However, the morphology of two kinds of colonies was different on Columbia blood agar plates. Δ*flpL* had smooth colony edges, but WT colony had untidy edges with fissures.

**Fig. 5.**
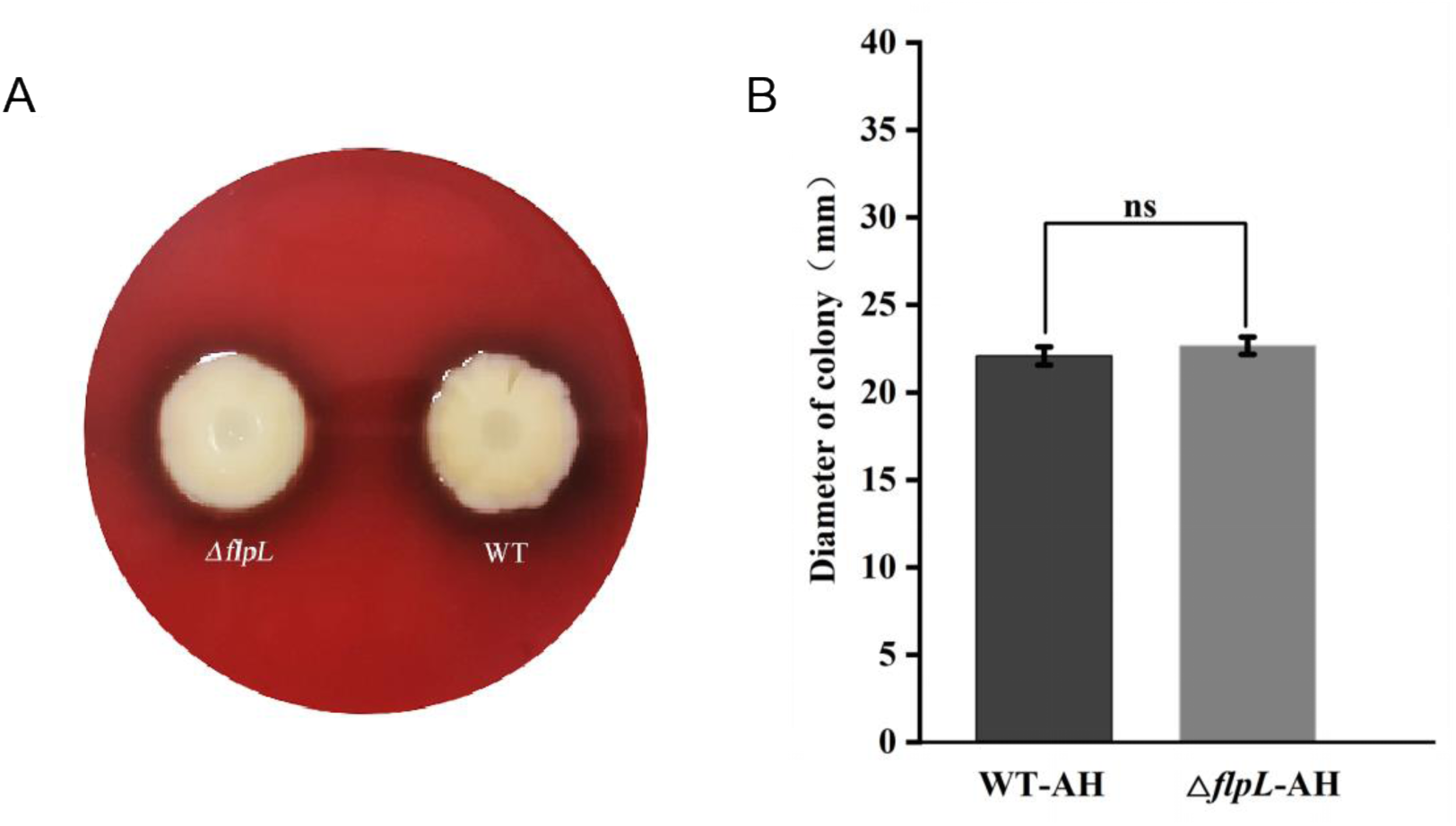
Comparison of haemolytic activity of Δ*flpL* and WT (ns represents not significant (*P* > 0.05), *means *P* < 0.05, **means *P* < 0.01). (A): Hemolytic activity assay of Δ*flpL* and WT. (B): Comparison of colony diameters between WT and Δ*flpL* bacterial strains.

### 3.6 Biofilm formation test

Through the 96-well plates method, the microplate reader detected OD_575_ values of 0.432 ± 0.005 for Δ*flpL* and 0.632 ± 0.011 for WT (Fig. 6). These results indicated that the *flpL* gene deletion reduced the biofilm formation capacity of *A. hydrophila*.

**Fig. 6.**
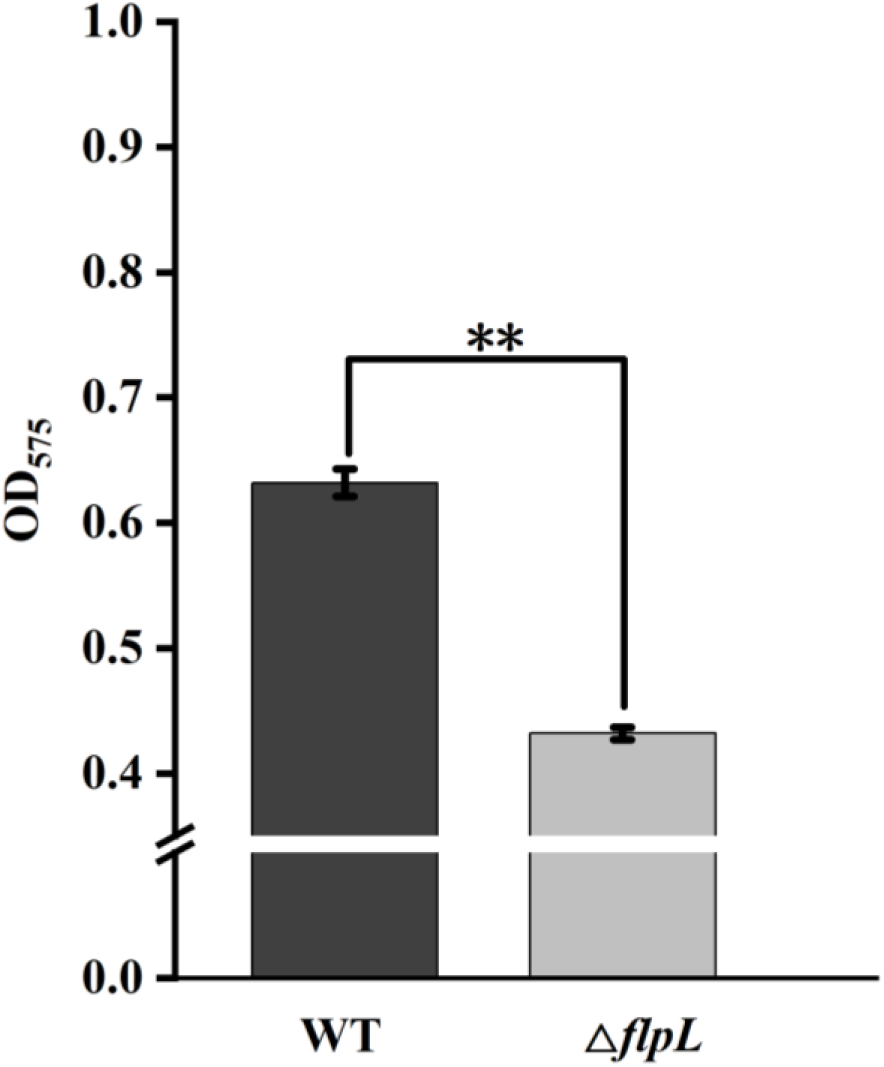
Biofilm formation of WT and Δ*flpL* strains (**means *P* < 0.01).

**Fig. 7.**
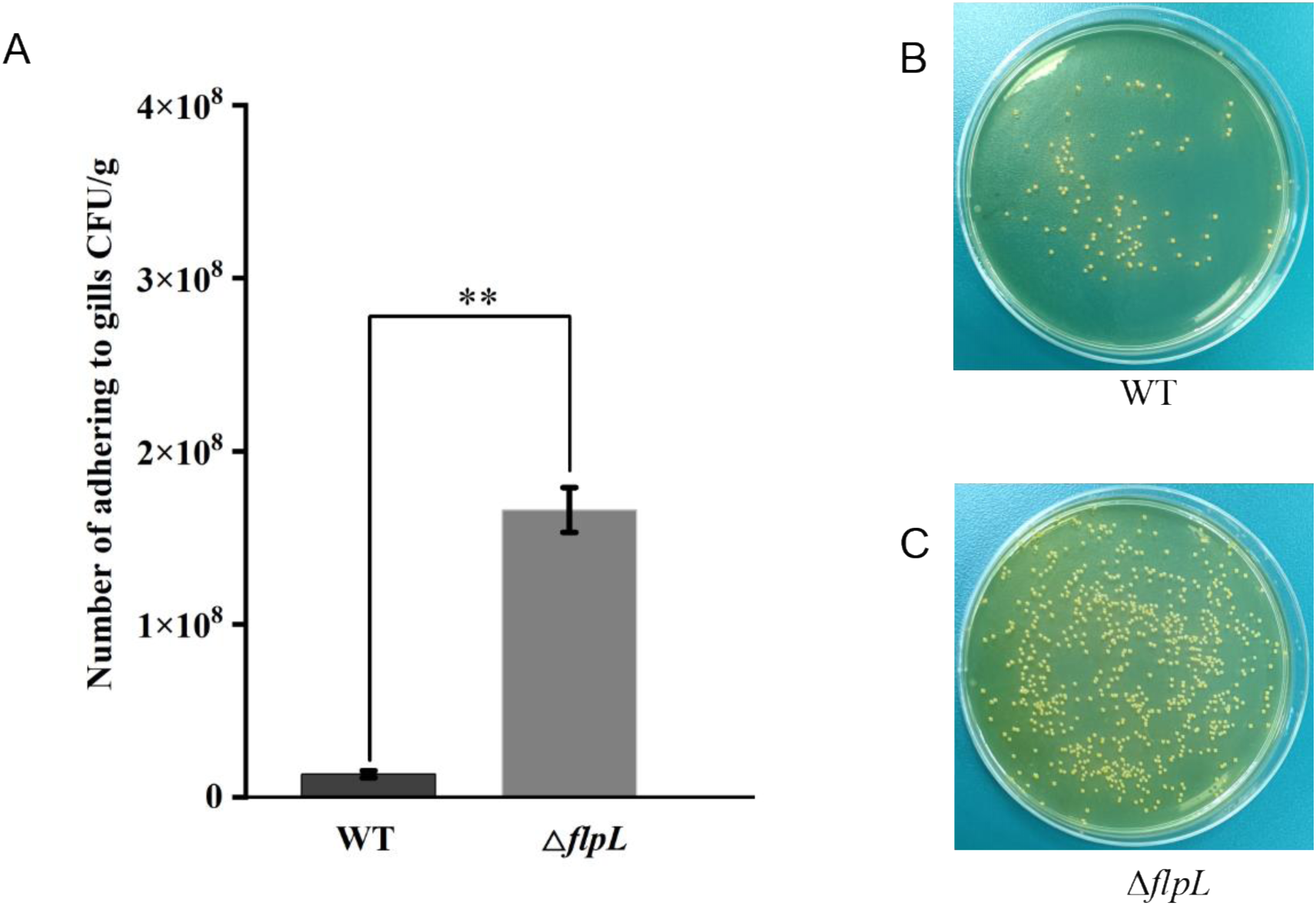
The adhesion levels between WT and Δ*flpL* (**means *P* < 0.01). (A): The quantity of bacteria attaching to the gills at 2 hours. (B): WT was coated following a 500-fold dilution. (C): Δ*flpL* was coated following a 500-fold dilution.

**Fig. 8.**
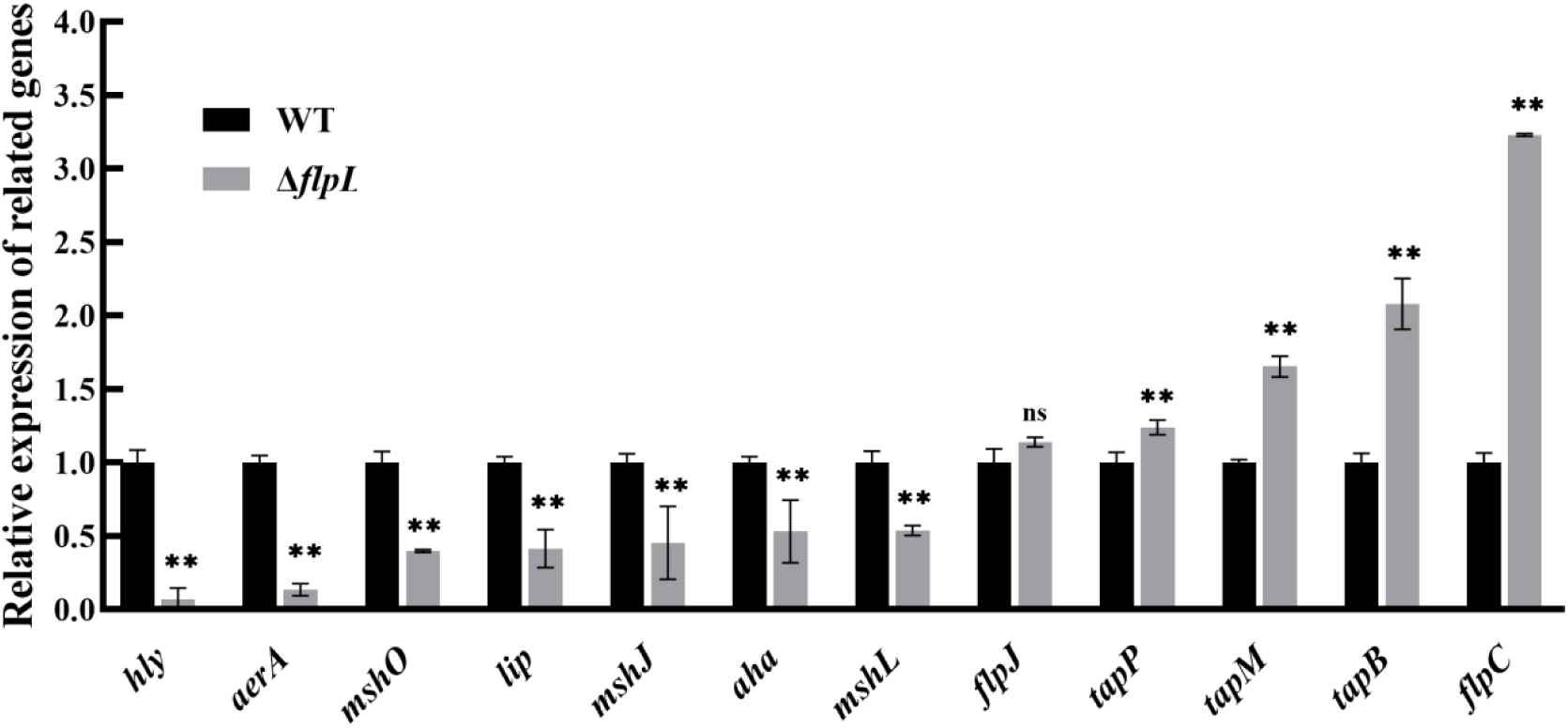
RT-qPCR result (ns represents no significance (*P* > 0.05), *means *P* < 0.05, **means *P* < 0.01).

### 3.7 Adhesion capacity

In our previous study, we constructed the standard curve that delineates the relationship between the quantity of *A. hydrophila* and the Ct value (Xiong *et al*, 2024). Based on the standard curve and equation (*y* = − 3.501*x* + 38.31), the adhesion levels of WT and Δ*flpL* were determined to be 1.32 × 10^7^ CFU/g and 1.66 × 10^8^ CFU/g, respectively. The adhesion capacity of the *flpL* gene was significantly enhanced, increasing 12.6-fold relative to WT. The Δ*flpL* group exhibited a notably higher number of bacterial colonies on RS solid medium compared to the WT group.

### 3.8 virulence gene expression

The expression levels of several virulence genes were analyzed in WT and Δ*flpL*. As illustrated in Fig. 9, *flpC* related Flp type IV pili and Tap type IV pili genes including *tapM*, *tapB*, *tapP*, showed significantly up-regulated (*P* < 0.05). Conversely, the genes associated with MSHA type IV pili, *lip*, *aha*, *aerA* and *hly*, exhibited notable down-regulation in Δ*flpL*.

## 3. Discussion

Flp pili, similar to other two types of T4P pili, greatly contribute to bacterial adherence, virulence, and capacity of biofilm formation (Long *et al*, 2024; Lu *et al*, 2024a). The *flpL* gene is a part of the T4P pili gene family. However, its function and the molecular mechanism in the virulence of *A. hydrophila* still unclear. This study makes the first attempt to elucidate the function of the *flpL* gene on virulence. The result demonstrated that the LD_50_ of Δ*flpL* was 4.87-fold higher than that of WT in Crucian carp, indicating the *flpL* gene is important for virulence of *A. hydrophila*. This finding aligns with study in *Actinobacillus pleuropneumoniae* (Li *et al*, 2019), *Haemophilus ducreyi* (Spinola *et al*, 2003), *Pasteurella multocida* (Fuller *et al*, 2000). Several studies have reported that unlike MSHA and Tap pili, the Flp pili appear to have a limited contribution to pathogenicity (Hadi *et al*, 2012; Ellison *et al*, 2022). However, in the study, the decrease in pathogenicity of the *flpL* gene was more significant than the deletion of genes related MSHA and Tap pili (Long *et al*, 2024; Lu *et al*, 2024a).

To investigated the mechanisms for the reduced pathogenicity in Δ*flpL*, biological characteristics associated with bacterial virulence were analyzed, including growth ability, motility, hemolytic activity, biofilm formation ability and cell adhesion. The results demonstrated that detection the *flpL* gene did not influence the growth capacity. The influence of the *flpL* gene on growth capacity was in accordance with previous studies in other type IV pili genes, such as *flp* gene in *Actinobacillus pleuropneumoniae* (Li *et al*, 2019) and *mshQ* gene in *A. hydrophila* (Qin *et al*, 2014). There is no significant difference observed in swarming capacity. The enhanced of swimming capacity in Δ*flpL* indicated that the *flpL* gene could affect *A. hydrophila* motility, which was consistent with some finding in previous studies that Flp pili gene deletions would affect bacterial motility (Carvia-Hermoso *et al*, 2024; Zatakia *et al*, 2014). Hemolysin is a one of the most critical virulence factors that plays critical role in bacterial pathogenicity (Zhao *et al*, 2024). The hemolytic activity of Δ*flpL* did not show a significant difference compared to that of WT, and both were β-hemolytic. However, the morphology of Δ*flpL* and WT strains had obvious difference. The edges of Δ*flpL* colonies are smooth, while the edges of WT colonies show a jagged shape. Previous studies have shown that the formation of folded and jagged colonies is linked to the biofilm formation capacity (Weissman *et al*, 2020). Therefore, we analyzed the difference of biofilm formation capacity between WT and Δ*flpL.* The findings demonstrated that a substantial reduction in biofilm formation occurred in Δ*flpL* compared with WT, indicating that the smooth edges of Δ*flpL* colonies on Columbia blood agar plates might be linked to the reduced biofilm formation capacity. Consistently, a previous study in *Xylella fastidiosa* also found that the type IV pili are essential for both motility and biofilm formation (Li *et al*, 2007). In addition, type IV pili play a key role in the adhesion processes of pathogenic gram-negative bacteria (Piepenbrink *et al*, 2014). Adhesion to host cells, which is the initial step of infection, heavily relies on the presence of pili. Studies have demonstrated that pili influence the adhesion capacity of *Aeromonas schubertii* (Piepenbrink *et al*, 2014), *Pseudomonas aeruginosa* (Beaussart *et al*, 2014), *Sulfolobus acidocaldarius* (Charles-Orszag *et al*, 2023). The present study demonstrated a significant enhance in the adhesion capacity of Δ*flpL*. The result was different from other previous studies on the deletion of genes related to the type IV pili (Qin *et al*, 2014; Li *et al*, 2023a). To sum up, Δ*flpL* exhibited a significant decrease in virulence for Crucian carp through reducing biofilm formation capacity, though swimming capacity and adhesion were enhanced.

To investigate the molecular mechanism of Δ*flpL* on pathogenicity, the expression levels of several critical virulence genes were analyzed, and cross-talks between *flpL* and other type virulence genes were found. The type IV pili is widely distributed fibers on bacterial surfaces that participate in diverse physiological behavior, including virulence, biofilm formation, motility and protein secretion (Ellison *et al*, 2022). The Δ*flpL* may affect the expression levels of other genes belonging to the same type IV pili genes. It is possible that the existence of overlapping ORF regions between different virulence factors in the bacterial genome, the deletion of one virulence factor using homologous recombination also impacts on the expression levels of other virulence genes in the same gene cluster. The current study found that the expression levels of *flpC*, *tapM*, *tapB* and *tapP* genes showed significant increase. The appearances of up-regulated expression of these genes might contribute to the enhanced swimming ability and increased adhesion capacity. The MSHA type IV pili primarily contribute significantly to biofilm formation (Hadi *et al*, 2012). The genes of MSHA type IV pili were down-regulated, which may explain the decrease in biofilm formation. Lipoprotein servers as a critical bacterial secretion system that plays a significant role in pathogenesis (Allaoui *et al*, 1992). The *aha* gene is critical for adhesion and pathogenicity (Song *et al*, 2019). Aerolysin, an exotoxin encoded by *aerA* gene, is highly cytotoxic, hemolytic, and enterotoxicity (Howard *et al*, 1987), playing an critical role in the virulence of *A. hydrophila* (Singh *et al*, 2008). The *hly* gene related to bacterial virulence, encodes pore-forming cytolysin listeriolysin. In the research, we speculate that the reduced expression of *lip*, *aha*, *aerA* and *hly* genes might be essential factors contributing to the decreased pathogenicity of the Δ*flpL* strain. However, the specific signaling pathway underlying this requires further investigation.

## 4. Conclusion

In conclusion, the first genetically stable *flpL* gene of *A. hydrophila* knockout strain was successfully constructed in this study. Our findings demonstrated that the Δ*flpL* exhibits significantly decreased pathogenicity for Crucian carp. This may be result from the decreased biofilm formation capacity and decreased expression levels of specific virulence-associated genes. Moreover, the significant increases in swimming capacity and adhesion may be related to the up-regulation of *flpC, tapM, tapB* and *tapP* genes. Motility and adhesion are strongly associated with the rate of bacterial infections. These results underscore the pivotal role of the *flpL* gene in pathogenicity, adhesion, motility, and biofilm formation capacity. This study will provide theoretical evidence supporting the potential development of Δ*flpL* as a live attenuated vaccine against *A. hydrophila* infections.

## 5. CRediT authorship contribution statement

**Hanyang Jiao**: Conceptualization, Methodology, Writing-Original Draft, Preparation, Writing-Review & Editing. **Kemei Liu**: Investigation, Methodology, Software. **ZiYi Liao**: Methodology, Software. **Adeeba Naseer**: Resources, Software. **Hua Ye**: Writing-Review & Editing. **Hao Xu**: Writing-Review & Editing. **Yun Li**: Writing-Review & Editing. **Yongyao Yu**: Writing-Review & Editing. **HuiQing Mei**: Writing-Review & Editing. **Ronghua Wu**: Conceptualization, Methodology, Validation, Writing-Review & Editing.

## 7. Data availability

The data that has been used is confidential.

## 8. Acknowledgements

This study was funded by National Natural Science Foundation of China (32102831).

## References

1. Allaoui A, Sansonetti P & Parsot C (1992) Mxij, a Lipoprotein Involved in Secretion of Shigella Ipa Invasins, Is Homologous to Yscj, a Secretion Factor of the Yersinia Yop Proteins. J Bacteriol 174: 7661–7669

2. Aly SM, Eissa AE, Abdel-Razek N & El-Ramlawy AO (2023) Chitosan nanoparticles and green synthesized silver nanoparticles as novel alternatives to antibiotics for preventing *A. hydrophila subsp. hydrophila* infection in Nile tilapia, Oreochromis niloticus. Int J Vet Sci Med 11: 38–54

3. Beaussart A, Baker AE, Kuchma SL, El-Kirat-Chatel S, O’Toole GA & Dufrene YF (2014) Nanoscale Adhesion Forces of *Pseudomonas aeruginosa* Type IV Pili. ACS Nano 8: 10723–10733

4. Boyd JM, Dacanay A, Knickle LC, Touhami A, Brown LL, Jericho MH, Johnson SC & Reith M (2008) Contribution of type IV pili to the virulence of *Aeromonas salmonicida* subsp salmonicida in Atlantic salmon (Salmo salar l.). Infect Immun 76: 1445–1455

5. Carvia-Hermoso C, Cuellar V, Bernabeu-Roda LM, van Dillewijn P & Soto MJ (2024) *Sinorhizobium meliloti* GR4 Produces Chromosomal– and pSymA-Encoded Type IVc Pili That Influence the Interaction with Alfalfa Plants. Plants-Basel 13: 628

6. Charles-Orszag A, van Wolferen M, Lord SJ, Albers S-V & Mullins RD (2023) Sulfolobus acidocaldarius adhesion pili power twitching motility in the absence of a dedicated retraction ATPase. bioRxiv

7. Chi Y, Jiao H, Ran J, Xiong C, Wei J, Ozdemir E & Wu R (2023) Construction and efficacy of Aeromonas veronii mutant Δhcp as a live attenuated vaccine for the largemouth bass (*Micropterus salmoides*). Fish & Shellfish Immunology 136: 108694

8. Ellison CK, Whitfield GB & Brun YV (2022) Type IV Pili: dynamic bacterial nanomachines. FEMS Microbiology Reviews 46: fuab053

9. Fang H-M, Ge R & Sin YM (2004) Cloning, characterisation and expression of *Aeromonas hydrophila* major adhesin. Fish & Shellfish Immunology 16: 645–658

10. Feng C, Liu X, Hu N, Tang Y, Feng M & Zhou Z (2022) Aeromonas hydrophila Ssp1: A secretory serine protease that disrupts tight junction integrity and is essential for host infection. Fish Shellfish Immunol 127: 530–541

11. Fernandez-Bravo A & Figueras MJ (2020) An Update on the Genus *Aeromonas*: Taxonomy, Epidemiology, and Pathogenicity. Microorganisms 8: 129

12. Finney DJ (1985) The median lethal dose and its estimation. Arch Toxicol 56: 215– 218

13. Fuller TE, Kennedy MJ & Lowery DE (2000) Identification of *Pasteurella multocida* virulence genes in a septicemic mouse model using signature-tagged mutagenesis. Microb Pathog 29: 25–38

14. Ganger MT, Dietz GD, Headley P & Ewing SJ (2020) Application of the common base method to regression and analysis of covariance (ANCOVA) in qPCR experiments and subsequent relative expression calculation. BMC Bioinformatics 21: 423

15. Gao Y, Tang X, Sheng X, Xing J & Zhan W (2016) Antigen uptake and expression of antigen presentation-related immune genes in flounder (*Paralichthys olivaceus*) after vaccination with an inactivated *Edwardsiella tarda* immersion vaccine, following hyperosmotic treatment. Fish Shellfish Immunol 55: 274–280

16. Hadi N, Yang Q, Barnett TC, Tabei SMB, Kirov SM & Shaw JG (2012) Bundle-Forming Pilus Locus of *Aeromonas veronii* bv. Sobria. Infect Immun 80: 1351– 1360

17. Howard SP, Garland WJ, Green MJ & Buckley JT (1987) Nucleotide sequence of the gene for the hole-forming toxin aerolysin of Aeromonas hydrophila. J Bacteriol 169: 2869–2871

18. Ji Y, Li J, Qin Z, Li A, Gu Z, Liu X, Lin L & Zhou Y (2015) Contribution of nuclease to the pathogenesis of *Aeromonas hydrophila*. Virulence 6: 515–522

19. Khalil W, Gantois C, Lemnos L, Salle L & Salle H (2021) *Aeromonas hydrophila* Is a Deceptive Pathogen Requiring Reconsideration of Antibiotic Prophylaxis. Surg Infect 22: 987–988

20. Li J, Ma S, Li Z, Yu W, Zhou P, Ye X, Islam MS, Zhang Y-A, Zhou Y & Li J (2021) Construction and Characterization of an Aeromonas hydrophila Multi-Gene Deletion Strain and Evaluation of Its Potential as a Live-Attenuated Vaccine in Grass Carp. Vaccines 9: 451

21. Li T, Zhang Q, Wang R, Zhang S, Pei J, Li Y, Li L & Zhou R (2019) The roles of *flp1* and *tadD* in *Actinobacillus pleuropneumoniae* pilus biosynthesis and pathogenicity. Microbial Pathogenesis 126: 310–317

22. Li Y, Han S, Wang Y, Qin M, Lu C, Ma Y, Yang W, Liu J, Xia X & Wang H (2023a) Autoinducer-2 promotes adherence of *Aeromonas veronii* through facilitating the expression of MSHA type IV pili genes mediated by c-di-GMP. Appl Environ Microbiol 89

23. Li Y, Hao G, Galvani CD, Meng Y, De la Fuente L, Hoch HC & Burr TJ (2007) Type I and type IV pili of *Xylella fastidiosa* affect twitching motility, biofilm formation and cell-cell aggregation. Microbiology-(UK) 153: 719–726

24. Li Y, Wei W, Wu J, Liu S, Ren Y, Huang X, Chen D, Geng Y & Ouyang P (2023b) Study a natural co-infection case of Largemouth bass ranavirus, Aeromonas vickert, and Aeromonas hydrophila in Micropterus salmoides. Isr J Aquac-Bamidgeh 75

25. Liang Y, Zhao H, Li Y, Gao F, Qiu J, Liu Z & Li Q (2024) Joint effects about antibiotics combined using with antibiotics or phytochemicals on Aeromonas hydrophila. Mar Environ Res 199: 106594

26. Liu Y, Wu Y, Srinivasan R, Liu Z, Wang Y, Zhang L & Lin X (2022) The protective efficacy of forty outer membrane proteins based DNA vaccines against Aeromonas hydrophila in zebrafish. Aquacult Rep 27: 101381

27. Long R, Wei J, Xiong C, Wang B, Lu J, Ye H, Li Y, Yu Y, Lin L & Wu R (2024) The role and function mechanism of tapP in modulating the virulence of Aeromonas hydrophila. Aquaculture 591: 741104

28. Lu J, Wei J, Liu K, Wang B, Zhang L, Yu Y, Li Y, Ye H, Li H & Wu R (2024a) MshK mutation reduces pathogenicity of Aeromonas veronii by modulating swimming ability, biofilm formation capacity, pili structure and virulence gene expression. Aquaculture 593: 741337

29. Lu J, Xiong C, Wei J, Xiong C, Long R, Yu Y, Ye H, Ozdemir E, Li Y & Wu R (2024b) The role and molecular mechanism of flgK gene in biological properties, pathogenicity and virulence genes expression of Aeromonas hydrophila. International Journal of Biological Macromolecules 258: 129082

30. Lukaszczyk M, Pradhan B & Remaut H (2019) The Biosynthesis and Structures of Bacterial Pili. In Bacterial Cell Walls and Membranes, Kuhn A (ed) pp 369–413. Cham: Springer International Publishing

31. Mansour A, Mahfouz NB, Husien MM & El-Magd MA (2019) MOLECULAR IDENTIFICATION OF *Aeromonas hydrophila* STRAINS RECOVERED FROM KAFRELSHEIKH FISH FARMS. Slov Vet Res 56: 201–208

32. Mohd-Aris A, Muhamad-Sofie MHN, Zamri-Saad M, Daud HM & Ina-Salwany MY (2019) Live vaccines against bacterial fish diseases: A review. Vet World 12: 1806–1815

33. Piepenbrink KH, Maldarelli GA, de la Pena CFM, Mulvey GL, Snyder GA, De Masi L, von Rosenvinge EC, Guenther S, Armstrong GD, Donnenberg MS, et al (2014) Structure of *Clostridium difficile* PilJ Exhibits Unprecedented Divergence from Known Type IV Pilins. J Biol Chem 289: 4334–4345

34. Qin YX, Yan QP, Mao XX, Chen Z & Su YQ (2014) Role of MshQ in MSHA pili biosynthesis and biofilm formation of *Aeromonas hydrophila*. Genet Mol Res 13: 8982–8996

35. Rasmussen-Ivey CR, Figueras MJ, McGarey D & Liles MR (2016) Virulence Factors of Aeromonas hydrophila: In the Wake of Reclassification. Front Microbiol 7

36. Sarkar P, Issac PK, Raju SV, Elumalai P, Arshad A & Arockiaraj J (2021) Pathogenic bacterial toxins and virulence influences in cultivable fish. Aquaculture Research 52: 2361–2376

37. Singh V, Rathore G, Kapoor D, Mishra BN & Lakra WS (2008) Detection of aerolysin gene in *Aeromonas hydrophila* isolated from fish and pond water. Indian J Microbiol 48: 453–458

38. Song H-C, Kang Y-H, Zhang D-X, Chen L, Qian A-D, Shan X-F & Li Y (2019) Great effect of porin(aha) in bacterial adhesion and virulence regulation in *Aeromonas veronii*. Microbial Pathogenesis 126: 269–278

39. Spinola SM, Fortney KR, Katz BP, Latimer JL, Mock JR, Vakevainen M & Hansen EJ (2003) *Haemophilus ducreyi* requires an intact *flp* gene cluster for virulence in humans. Infect Immun 71: 7178–7182

40. Stepanović S, Vuković D, Dakić I, Savić B & Švabić-Vlahović M (2000) A modified microtiter-plate test for quantification of staphylococcal biofilm formation. Journal of Microbiological Methods 40: 175–179

41. Suresh K & Pillai D (2023) Prevalence and characterization of virulence-associated genes and antimicrobial resistance in *Aeromonas hydrophila* from freshwater finfish farms in Andhra Pradesh, India. Biologia 78: 2931–2939

42. Tomich M, Planet PJ & Figurski DH (2007) The *tad* locus:: postcards from the widespread colonization island. Nat Rev Microbiol 5: 363–375

43. Weissman Z, Pinsky M, Wolfgeher DJ, Kron SJ, Truman AW & Kornitzer D (2020) Genetic analysis of Hsp70 phosphorylation sites reveals a role in *Candida albicans* cell and colony morphogenesis. BBA-Proteins Proteomics 1868: 140135

44. Wu R, Shen J, Lai X, He T & Li Y (2020) Development of monoclonal antibodies against serum immunoglobulins from gibel carp (*Carassius auratus gibelio*) and their applications in serodiagnosis of inapparent infection and evaluation of vaccination strategies. Fish Shellfish Immunol 96: 69–77

45. Xiong C, Xiong C, Lu J, Long R, Jiao H, Li Y, Wang B, Lin Y, Ye H, Lin L, et al (2024) flgL mutation reduces pathogenicity of Aeromonas hydrophila by negatively regulating swimming ability, biofilm forming ability, adherence and virulence gene expression. International Journal of Biological Macromolecules 261: 129676

46. Yang S, Mkingule I, Liu L, Chen W, Yuan X, Ma Z, Liang L, Qian S, Huang M & Fei H (2023) Protective efficacy evaluation of immunogenic protein AHA_3793 of *Aeromonas hydrophila* as vaccine candidate for largemouth bass *Micropterus salmoides*. J Oceanol Limnol 41: 392–400

47. Yi Y, Zhang H, An Y & Chen Z (2024) A Live Attenuated H1N1 Influenza Vaccine Based on the Mutated M Gene. Vaccines 12: 725

48. Zatakia HM, Nelson CE, Syed UJ & Scharf BE (2014) ExpR Coordinates the Expression of Symbiotically Important, Bundle-Forming Flp Pili with Quorum Sensing in Sinorhizobium meliloti. Applied and Environmental Microbiology 80: 2429–2439

49. Zhang M, Zhang T, He Y, Cui H, Li H, Xu Z, Wang X, Liu Y, Li H, Zhao X, et al (2023) Immunogenicity and protective efficacy of OmpA subunit vaccine against *Aeromonas hydrophila* infection in *Megalobrama amblycephala*: An effective alternative to the inactivated vaccine. Front Immunol 14: 1133742

50. Zhang Z, Liu G, Ma R, Qi X, Wang G, Zhu B & Ling F (2020) The immunoprotective effect of whole-cell lysed inactivated vaccine with SWCNT as a carrier against *Aeromonas hydrophila* infection in grass carp. Fish Shellfish Immunol 97: 336– 343

51. Zhao X, He J, Liu J, Deng H, Pan Y & Ye S (2024) Arbutin interacts with *Vibrio harveyi* hemolysin to alleviate damage from associated infection. Aquaculture 584: 740633

